# Ceftolozane-tazobactam pharmacokinetics during extracorporeal membrane oxygenation in a lung transplant recipient

**DOI:** 10.1101/438671

**Authors:** Fabio Arena, Luca Marchetti, Lucia Henrici De Angelis, Enivarco Maglioni, Martina Contorni, Maria Iris Cassetta, Andrea Novelli, Gian Maria Rossolini

## Abstract

Ceftolozane-tazobactam pharmacokinetics during extracorporeal membrane oxygenation (ECMO) has not been previously studied. In this work we report on the ceftolozane and tazobactam plasmatic levels in a lung transplant recipient during ECMO, treated with ceftolozane-tazobactam (2g/1g, intravenously every 8 h, 1 h infusion) for a *Pseudomonas aeruginosa* pulmonary infection. Ceftolozane Cmax and Cmin, monitored during 96 hrs, remained above 60 and 20 μg/mL, respectively, with optimal drug exposure (100% %*T*_MIC_). Tazobactam levels were above 1.9 μg/mL.

---

Ceftolozane-tazobactam (Merck Sharp & Dohme Corp. NJ, USA) is a recently introduced β-lactam plus β-lactam inhibitor combination targeting Gram-negative pathogens, including multi-drug resistant (MDR) *Enterobacterales* and *Pseudomonas aeruginosa*, with the exclusion of carbapenemase-producing strains. At the registered dose of 1.5 g (1 g/0.5 g) every 8 hours, ceftolozane-tazobactam is currently approved for the therapy of complicated urinary tract and intra-abdominal infections (in the latter case in association with metronidazole). (1, 2) Ceftolozane-tazobactam has been successfully used for treatment of lower respiratory tract infections (LRTI) caused by MDR *P. aeruginosa*, also in cystic fibrosis patients. (3–5) Results of a phase 3 safety and efficacy study of ceftolozane/tazobactam to treat ventilated nosocomial pneumonia (MK-7625A-008) will be available in the next future.

The PK/PD index of ceftolozane that best correlates with *in vivo* efficacy is the percentage of the time in which free plasma drug concentrations exceed the minimum inhibitory concentration of a given pathogen %*T*_MIC_). (6) The time above a threshold concentration has been determined to be the parameter that best predicts the efficacy of tazobactam in *in vitro* nonclinical models (threshold = 0.25 μg/ml, for isolates with high-β-lactamase expression). (7)

In healthy adult subjects, mean Cmax serum values of ceftolozane after 1 to 10 days of 1000 mg administration are around 110-120 μg/mL. (8) Considering a plasma-to-epithelial lining fluid penetration ratio of approximately 50%, a 3 g dose of ceftolozane-tazobactam is required to achieve a >90% probability of target attainment against pathogens with an MIC of ≤8 μg/mL, in patients with pneumonia and a normal renal function. (9) At this dosage, ceftolozane-tazobactam represents an attractive option for treatment of serious *P. aeruginosa* LRTI, including those occurring in patients undergoing lung transplantation.

In this latter category, the use of extracorporeal membrane oxygenation (ECMO) as a bridge to transplantation or support to lung function in case of severe lung injury is increasingly popular. (10) However, data regarding ceftolozane-tazobactam pharmacokinetics in critically ill patients during ECMO support are lacking.

Here we report on the ceftolozane and tazobactam plasmatic levels documented in a lung transplant recipient during ECMO, treated with ceftolozane-tazobactam for a *P. aeruginosa* infection.

In September 2017, a 42-year-old female patient, weighting 65 Kg (BSA 1,72 m^2^) was admitted to the Cardiothoracic Intensive Care Unit of Siena University Hospital (Italy) due to acute respiratory failure. She had a history of sarcoidosis with pulmonary fibrosis (Stage IV) and pulmonary emphysema with alpha-1 antitrypsin deficiency. The chest Xrays showed bilateral inflammatory infiltrates. The patient had a history of *P. aeruginosa* chronic respiratory infections and, basing on previous susceptibility testing data, an antibiotic therapy was started with vancomycin and meropenem. On day 2 after admission, following a worsening of hypoxia despite mechanical ventilation, a veno-venous ECMO (vvECMO) support was started according to the ELSO guidelines (www.elso.org). A PLS circuit system (MAQUET Holding B.V. & Co. KG., Rastatt, DE) with a pre-connected standard set consisting of a PLS-i oxygenator and a centrifugal pump (ROTAFLOW), both incorporated into a tubing set with tip-to-tip BIOLINE coating, was used. Blood and gas flows were adapted to maintain PaO_2_ >90 mm Hg, PCO_2_ < 40 mm Hg and pH 7.4, under protective ventilation.

The tracheal aspirate culture, performed on day 2 after admission, grew a carbapenem-resistant *P. aeruginosa* (Table 1), Based on susceptibility testing results, the antimicrobial therapy was modified to vancomycin, amikacin and ciprofloxacin. However, the culture of the tracheal aspirate, performed on hospital day 10, was still positive for *P. aeruginosa* >1 × 10^6^ CFU/mL with an identical antimicrobial susceptibility profile.

**Table 1.** Susceptibility profile of the *Pseudomonas aeruginosa* isolate obtained from tracheal aspirate culture.

| Antibiotic | Minimal inhibitory<br>concentration in µg/mL | Categorization* |
| --- | --- | --- |
| amikacin | 4 | susceptible |
| aztreonam | 16 | resistant |
| ceftazidime | 2 | susceptible |
| ciprofloxacin | 0.5 | susceptible |
| colistin | 0.5 | susceptible |
| gentamicin | 1 | susceptible |
| imipenem | 16 | resistant |
| meropenem | 16 | resistant |
| piperacillin/tazobactam | 32/4 | resistant |
| ceftolozane/tazobactam | 4/4 | susceptible |

\*interpretation according to EUCAST clinical breakpoints (v\_8.1). Susceptibility testing was performed by reference broth microdilution method (13).

On hospital day 13 the patient was subjected to bilateral lung transplantation. During surgery, the extracorporeal support was switched from veno-venous to veno-arterial (vaECMO) and then maintained for hemodynamic instability.

Immediately after lung transplantation, the antibiotic therapy was modified to ceftolozane/tazobactam 3 g intravenously every 8 h over 1 h infusion, ciprofloxacin 400 mg every 8 h and vancomycin in continuous infusion (vancomycin dose was adjusted basing on TDM). The patient had a normal renal function with creatinine clearance, estimated using the Cockcroft-Gault formula, above 90 mg/min (range 90-150 mg/min) (Figure 1).

**Figure 1.**
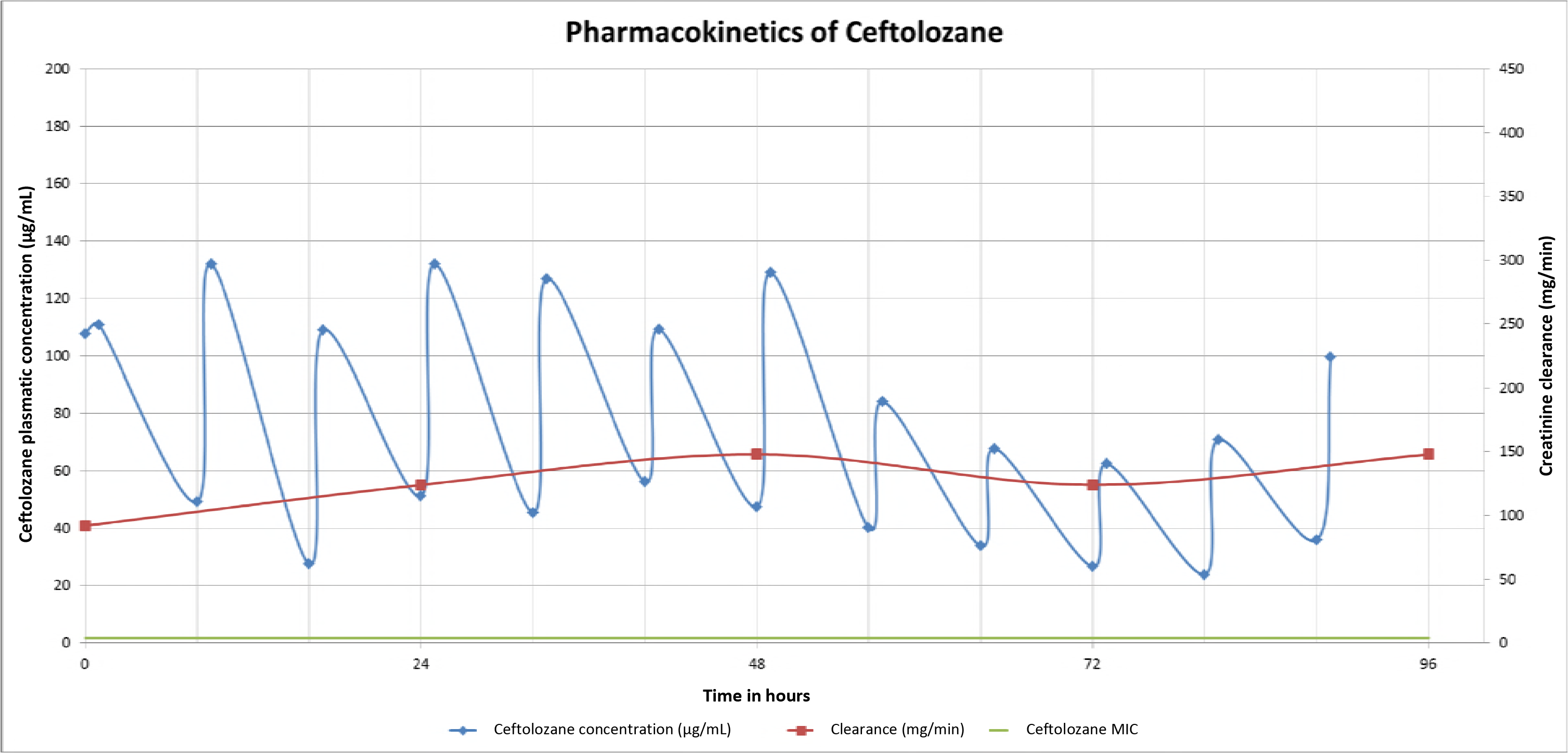
96 hrs monitoring ceftolozane plasmatic concentrations.

Ceftolozane and tazobactam plasma concentrations were monitored for 96 h following the first administration. Plasma samples were obtained ½ h after the end of each ceftolozane-tazobactam infusion and immediately before the beginning of the following dose (Cmax and Cmin respectively). The samples were centrifuged at 1,600 *g* for 10 min to yield at least 1 ml of plasma. Plasma samples were stored at −80° until the assay. Ceftolozane and tazobactam concentrations were quantified using a validated HPLC method at the Chemotherapy Laboratory of the Department of Health Sciences (University of Florence, Italy). (11)

In the following days the patient clinical conditions improved and, on day 3 after the lung transplant, the vaECMO was switched again to vvECMO.

In the 96 h of drug monitoring, the ceftolozane levels remained constantly above the isolate MIC. In the first 48 h, ceftolozane Cmax were persistently above 100 μg/mL. Later, from 48 to 96 h, there was a decline in Cmax and Cmin, which, however, remained above 60 and 20 μg/mL, respectively (Figure 1). Tazobactam concentration were above 1.9 μg/mL for all the monitored period.

ECMO was halted on day 8 after lung transplantation. The patient’s conditions eventually improved and, on day 15 after lung transplantation, the antimicrobial therapy was discontinued. No adverse events were attributed to the ceftolozane-tazobactam administration. Four consecutive tracheal aspirate cultural exams, performed in the last day of treatment and 3, 4, and 7 days after the antibiotics discontinuation, did not grow *P. aeruginosa* or any other pathogen.

Concluding, we documenting the success of ceftolozane-tazobactam treatment for a case of carbapenem-resistant *P. aeruginosa* LRTI in a lung transplant recipient subjected to ECMO support. TDM revealed that optimal drug exposure was achieved, with a ceftolozane concentration at 100% %*T*_MIC_. Of note, in this case, optimal Cmax and Cmin were obtained using a 3 g cefotolozane-tazobactam dose every 8 h, using a standard 1 h infusion time. During the TDM period we assisted to a decrease of ceftolozane/tazobactam Cmin and Cmax that could be attributed to the increase in the patient’s creatinine clearance values. In fact, it was previously demonstrated that renal function is the most significant variable influencing the PK of ceftolozane/tazobactam (12).

Our findings suggest that optimal ceftolozane-tazobactam PK parameters can be achieved in severely ill patients with normal renal function, requiring ECMO, without the need of dose or infusion time adjustment.

## Acknowledgments

This research received no specific grant from any funding agency in the public, commercial, or not-for-profit sectors. AN has provided consultancies/advisory services to and received research funding from Merck Sharp & Dohme Corp. GMR has provided speaker bureau and consultancies and received research grants from Merck Sharp & Dohme Corp.

